# Study of compatibility and determination of the suitable pollinizer for Argane tree (*Argania spinosa* L.)

**DOI:** 10.1101/2021.05.23.445304

**Authors:** Naima Ait Aabd, Abdelghani Tahiri, Abdelaziz Mimouni, Rachid Bouharroud

**Affiliations:** National Institute of Agronomic Research, CRRA Agadir, Morocco

**Keywords:** Argane tree, breeding program, pollinizers, self-incompatible, crossing diallel programs, compatibility

## Abstract

During the breeding program studies, the pollen donor parents (pollinizers) were observed to be characterized by a strong flowering rate and very low fruit set, even after hand pollination. Then the notion of pollinizers in the argane species was born, first mentioned, checked, documented and confirmed like other conventional tree species. Since the argane tree is completely self-incompatible, the presence of compatible pollinizers is necessary for the fruit set. Indeed, pollinizers need to be selected accurately based on the synchronization of bloom periods and compatibility to receiver trees (female). The pollinizer is essential in any breeding program and for new orchard plantations. The current study was conducted on 13 argane genotypes including two pollinizers. The flowering period, bloom phenology, floral structure and fruit set of crossed genotypes were monitored, illustrated and a season phenogram was established. The pollen viability and germination were also evaluated. In order to test compatibility, the hand pollination using two selected pollinizer pollens was compared to open pollination. Then, the compatibility system was monitored and evaluated through analysis of crossing diallel programs and through the index of self-incompatibility. The flowering periods are genotype depending and one to three blooms have been observed during the two years study (2018-2019) and the argane tree is a tristylous species (Mesostylous, brevistylous and longistylous flowers). The *in vitro* tests showed that the pollen originated from crossed genotypes were viable and able to germinate. The cross-compatibility rate depends on cross associations and it varies from 39 to 84 %. In fact, this study showed that the effect of pollen-parent (xenia) occurs in all fruit components of argane tree. It was observed, for the first time, that both compatible pollinizers and xenic effects of pollen on argane fruit have occurred. Artificial pollination is currently feasible for breeding programs and the screening of elite genotypes. Then the selected pollinizer is quite required for the development of argane tree cropping.

**Highlights:**

- Pollinizers in the argane species first mentioned
- Pollinizer is essential in any breeding program and for new orchard plantations
- Argane tree is self-incompatible
- Pollen-parent effect, (xenia) occurs in all fruit components of argane tree

## Introduction

The argane tree (*Argania spinosa* (L.) Skeels, previously *Argania sideroxylon* Roem. & Schult.) a unique representative of the sapotaceae family in Morocco, is one of the most important oil seed plants in the world. It was previously called *Sideroxylon spinosum*, then *Argania sideroxylon*. However, a species belonging to a related genus (*Sideroxylon marmulano* Banks). It is widely distributed in arid and sub arid of the Southwest region, and three relics populations of small sizes are in the North at Oued Grou (close to Rabat), Northeast at Beni Snassen and South at Guelmim (Ehrig, 1974; Prendergast and Walker, 1992) covering over 900,000 ha (HCEFLCD, 2012).

Several studies were carried out and characterized the great genetic diversity of the argane tree (Msanda 1993; El Mousadik and Petit, 1996; Ait Aabd et al., 2013; Ait Aabd et al., 2014; Yatrib et al., 2015; Ait Aabd et al., 2019) and its geographical distribution (from sea level to 1400-1500 meters altitude). Since there is an intensive demand at international market level, it is obvious that policy makers in Morocco pays accentuate efforts for the conservation of this endemic and endangered species. It then becomes urgent to think of a sustainable use of this resource as a preferred way to ensure its protection, by creating the National Agency for the Development of Oasis and Argane Ecosystems (ANDZOA). Also, the planting program of 50 000 Ha was developed and initiated in accordance with the new agricultural strategy “GREEN GENERATION 2020-2030”.

The argane tree is a perennial, monoecious tree with hermaphrodite flowers. The flowers are protogynous, the style emerging from the flower before anthesis. Several works have been published on the floral biology of argane trees, flowering periods, pollination and fruit maturation (Perrot, 1907; Nerd et al., 1998; Belmouden and Bani-Aameur, 1995; Benlahbil, 2003; Ait Aabd et al., 2019; Ajerrar et al., 2020). The pollination of the argane tree is allogamous. So, need a vector for pollination either by insects^15^ or by the wind. This vector is also called a pollinator and the difference between pollinator and pollinizer term used in this study is simply the tree that provides viable abundant compatible pollens to female flower. The pollinizer in this study refers to pollen donor. The knowledge in flowering, pollination and fertilization in the argane tree are still very limited. Many imprecisions and controversies persist, which justifies further research on the mode of reproduction in order to control the actions to be taken for the domestication and pollination strategies to improve fruit set in Argane orchards.

The synchronization of the flowering period, the availability of compatible pollen grains, or compatible trees, the presence of vectors to transfer pollen to the stigma, are essential to the success of fruit production. The argane tree is characterized by a high intensity of flowering, but the fruit set rate is very low, due to the massive and premature fall of the flowers at the beginning of bloom and no information was available to explain the reasons for this fall in the argane tree. For fruit trees (almond, olive, citrus…), the most important factor in fruit set and fertilization is compatibility (Ortega and Dicenta, 2004). Self-incompatibility or incomplete pollination of flowers is one of the main causes of fruit drop (Boöković and Tobutt 2001), it plays a major role in reducing yield for some fruit trees. Compatibility and incompatibility in the argane tree should be studied to have enough information for the improvement of fruit yield.

There are several methods to study the compatibility or incompatibility of different cultivars and to determine the suitable pollinizers (Ortega and Dicenta, 2004; Mousavi et al., 2014). These methods include controlled pollination which allows to estimate the performance of various cultivars in the orchard, so this method is recommended to determine the appropriate pollinizers (Rasouli et al., 2009). In order to improve the knowledge of the compatibility mechanisms of the argane tree, the compatibility was studied of selected argane genotypes with two pollinizers. The notion of pollinizers in the argane tree species has its origin from this study and the main aim is the confirmation of this finding through appropriate tests. Thus, the current study aimed to check if there are different flowering patterns among different genotypes after morphological characterization in the orchard, to test the cross and self-incompatibility of pollination of this monoecious species with hermaphrodite flowers, to describe the polymorphism of the argane tree flowers and finally to test if crosspollination leads to large fruit size than self-pollination (metaxenic effect).

## Materials and methods

### Orchard design

The experiments were conducted in argane orchard (2 Ha) of CRRA-Agadir in experimental farm Melk Zhar, Belfaa, located at 36 km from Agadir (30.0434N; −9.55635W; alt. 100 m). This orchard has been planted in 2010 as an experiment of argane tree domestication at a density of 8 × 6 m (**Fig. 1**). These trees are the elite individuals selected from the natural area distribution of argane. All trees were drip irrigated simultaneously and at the same frequency and dose.

**Fig. 1.**
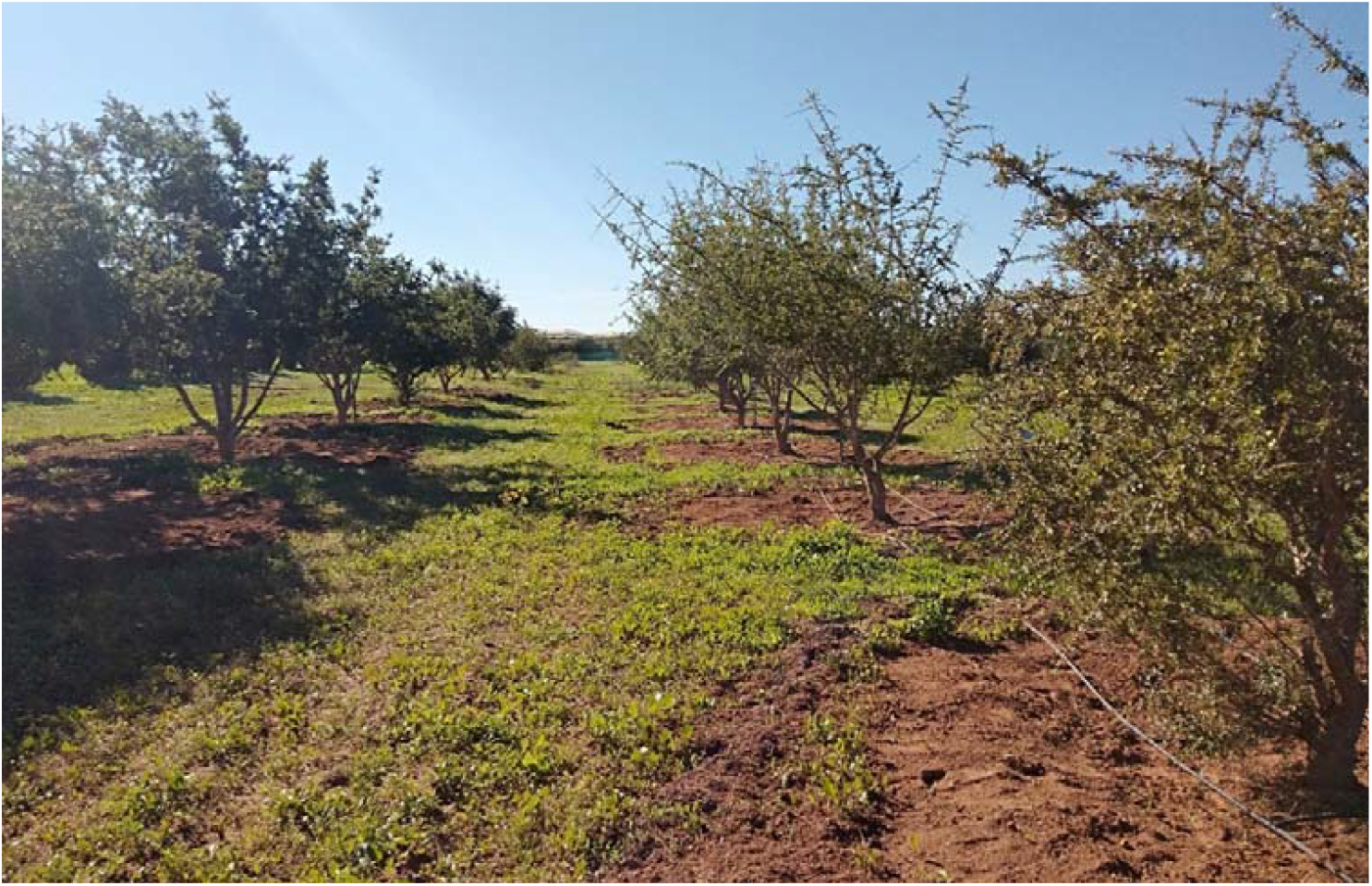
Argane tree orchard located at Melk Zhar experimental farm.

### Plant materials

This research was carried out during crop seasons 2018 and 2019. Two promising pollinizers genotypes labeled INRA-62 and INRA-142 as pollen donor parent was crossing with eleven genotypes of argane trees. The INRA-62 pollinizer was crossed to seven genotypes (INRA-139, INRA-135, INRA-109, INRA-98, INRA-79, INRA-65, INRA-49) and INRA-142 pollinizer was crossed to six trees (INRA-132, INRA-107, INRA-98, INRA-65, INRA-75, INRA-54). Morphological characteristics (height, trunk number, trunk collar diameter, number of the branch at a collar and the circumference of the tree), flowering and fruiting period, self-cross-compatibility and fruit set of the donor parent (or pollinizers) and eleven female parents (receptor pollen) were described in **Table 1, Fig. 2, 3** and **4**.

**Table 1.** Morphological characteristics of two pollinizers (INRA-142; INRA-62) and 11 receiver genotypes.

| Tree | Tree form | Tree height (m) | Number of trunk | Collar diameter (cm) | Number of branches at the collar | Tree circumference (m) |
| --- | --- | --- | --- | --- | --- | --- |
| **INRA-62** | draw up | 3.20 | 1 | 37 | 2 | 12.00 |
| **INRA-142** | draw up | 2.60 | 1 | 56 | 2 | 12.66 |
| **INRA-49** | draw up | 5.00 | 1 | 61 | 6 | 16.66 |
| **INRA-54** | draw up | 3.55 | 1 | 51 | 4 | 10.66 |
| **INRA-75** | Semi-weeping | 3.80 | 1 | 63 | 5 | 14.00 |
| **INRA-79** | draw up | 3.25 | 1 | 44 | 3 | 9.33 |
| **INRA-98** | draw up | 3.95 | 1 | 62 | 3 | 14.33 |
| **INRA-109** | draw up | 3.50 | 1 | 37 | 2 | 12.00 |
| **INRA-132** | draw up | 3.80 | 3 | 76 | 4 | 11.33 |
| **INRA-135** | draw up | 2.95 | 1 | 44 | 2 | 12.00 |
| **INRA-139** | draw up | 3.60 | 1 | 62 | 3 | 14.00 |
| **INRA-107** | draw up | 2.60 | 1 | 36 | 3 | 9.33 |
| **INRA-65** | draw up | 4.35 | 1 | 62 | 2 | 14.66 |

**Fig. 2.**
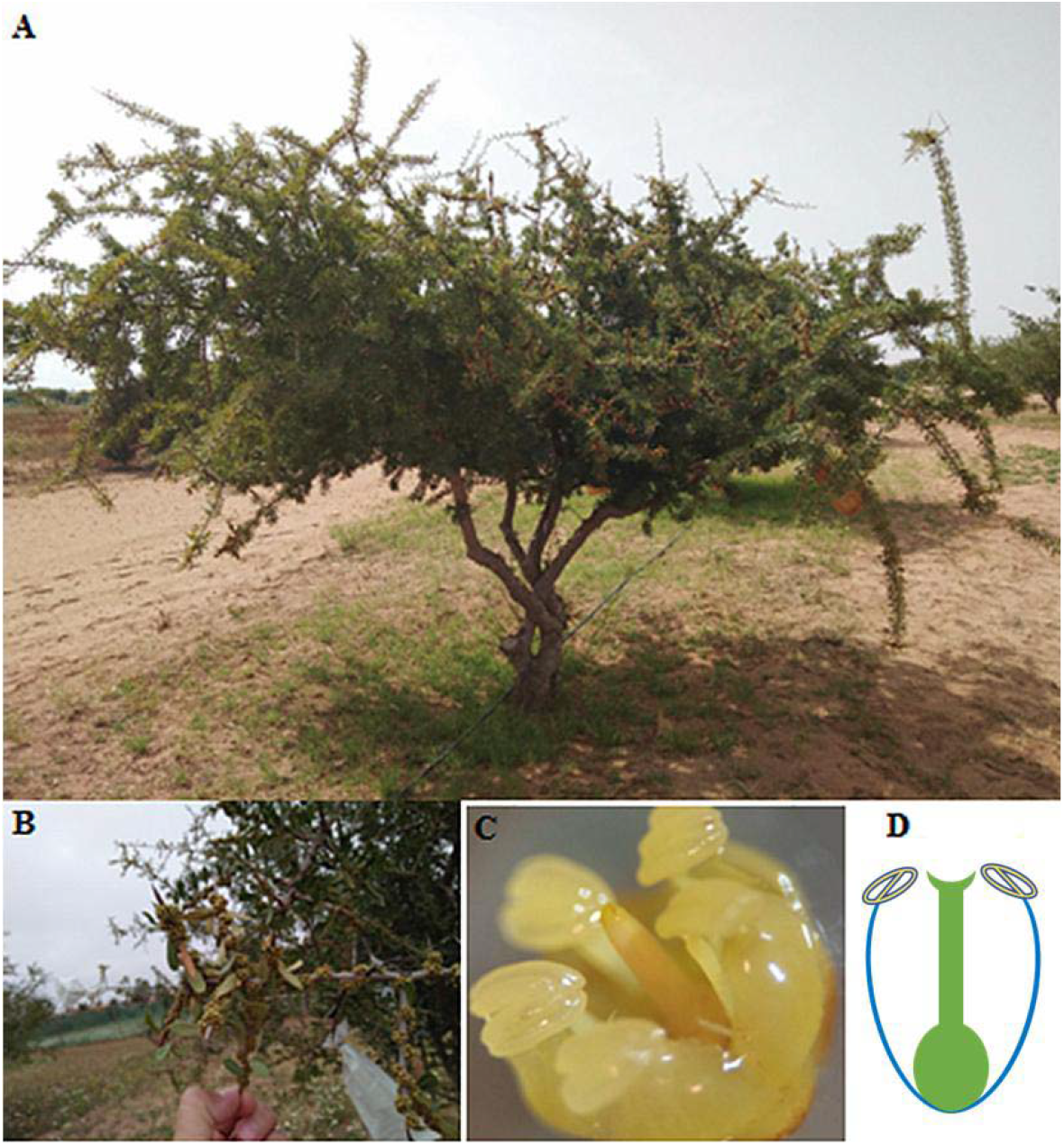
Morphological variability in domesticated pollinizer (A) and wild pollinizer (E): **A** (tree morphology of domesticated pollinizer; INRA-142), **B** (a large number of flower buds per branch), **C-D** (flower morphology of pollinizer), **E** (floral potential or a higher quantity of pollen of wild pollinizer)

**Fig. 3.**
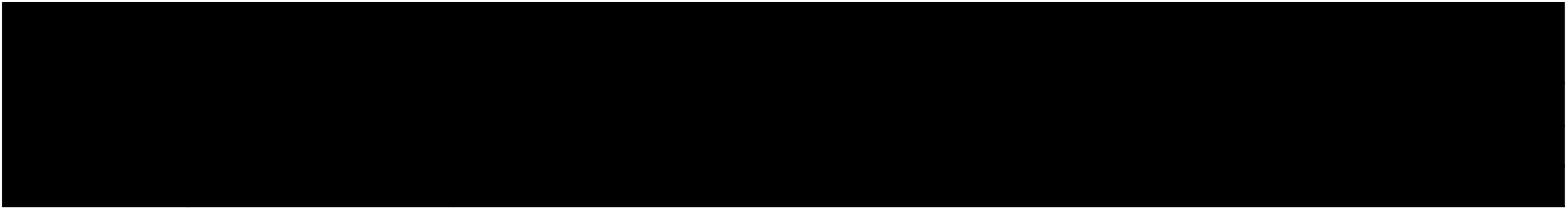
A flowering period of two argane pollinizers. Full bloom is showed in orange color.

**Fig. 4.**
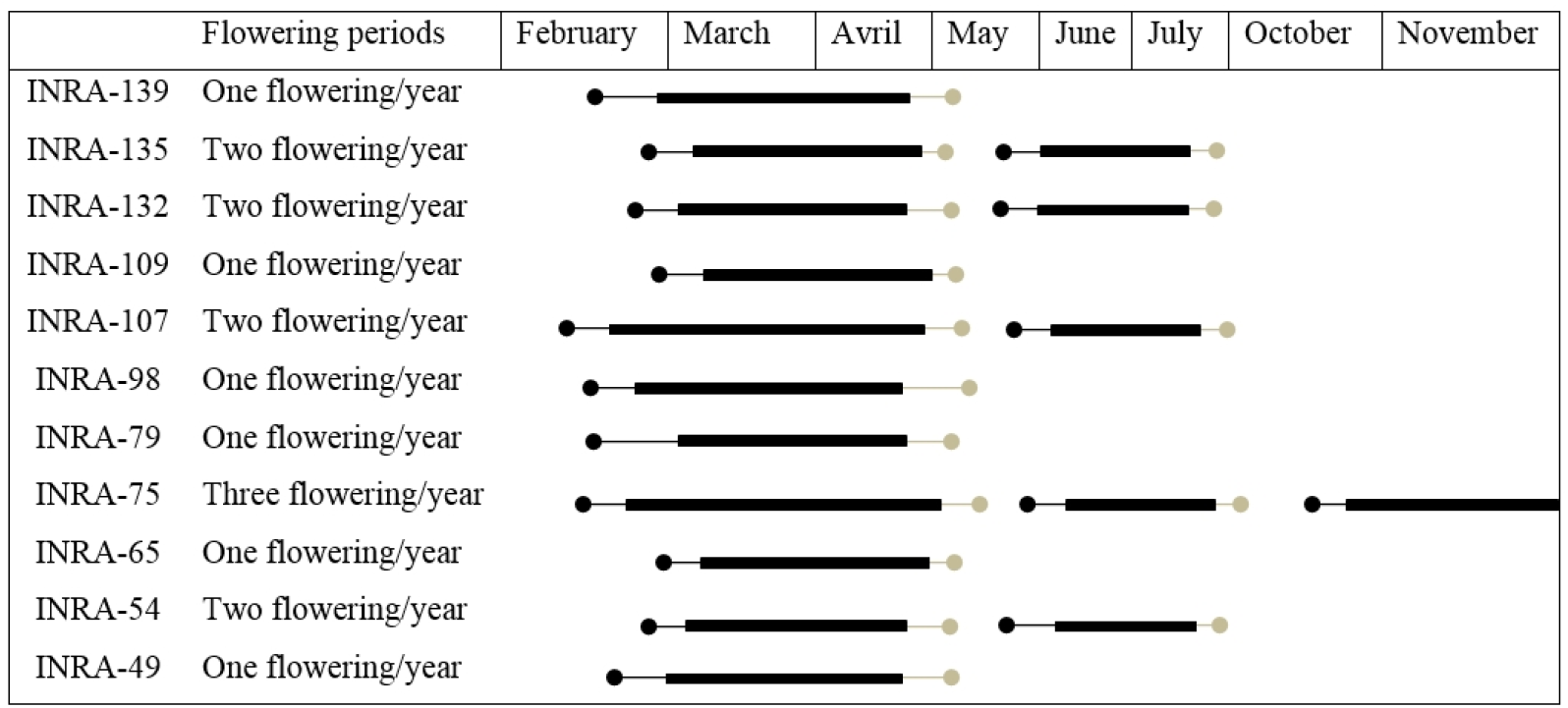
Diagram of flowering of the 11 argane genotypes crossed with pollinizers. Full bloom is showed in dark color.

### Flowering period and bloom phenology

The flowering period of parent’s trees was observed over the two consequent years (2018-2019), and stages were divided as follows: the beginning of bloom (first flower bud), full bloom (open flowers), and the end of bloom (total petals fall). The observations were conducted for each parent on three periods per year, from February to April (bloom once a year), and from June to July (bloom twice a year) and length of the year for a single genotype that blooms three times a year. Season phenograms of flowering were made for all trees evaluated in this study based on the collected data of the bloom period (**Fig. 3** and **Fig. 4**).

### Pollination

Pollination was done in March-April of 2018–2019. The flower buds were bagged before anthesis to prevent pollination with alien pollen and hand-pollinated with pollen collected from two pollinizer trees, after emasculation.

### The crossbreeding design and field pollination experiment

The flowers of eleven receiver trees were assigned randomly to one of three treatments: cross-pollination, self-pollination and no-pollination. Branch with a large number of flowers of donor and receiver parents were bagged before and after anthesis. In the morning, from 07:30 to 10:30 h, before anther dehiscence, the donor pollens were collected and flowers of the receiver parents were emasculated then pollinated. Thus, pollinated flowers were counted, labeled and bagged. To ensure pollination, flowers were pollinated again with donor pollen 24 h later. Open pollination was considered as a control treatment, although cross (hand-pollination) and self-pollination were used for the study of compatibility treatment. For each cross, 676 flowers were emasculated and pollinated. For self-pollination treatment, 1130 flowers were bagged without emasculation. The compatibility system was evaluated through analysis of crossing diallel programs and through the index of self-incompatibility (ISI). So, the ISI ratio was calculated between the percentage fruit set resulting from hand-self-pollination over that from hand-cross-pollination. As a result, the specie is considered as a self-compatible (SC) when ISI >0.25, and as a self-incompatible (SI) when ISI < 0.25 (Bawa, 1974). The fruit set for all types of pollination was recorded 30 days after the end of bloom (initial fruit set) and was expressed as a percentage.

### Floral structure polymorphism and study of effect of pollinizers and floral morphs on fruit set and fruit weight

To determine the floral polymorphisms and if the different morphs in the argane tree (different heights of stigmas and anthers) influence the fruit set and fruit weight, the floral structure was observed in the field and the laboratory. During the bloom (March-June / 2018), the flowers were collected at the late balloon stage, fixed in Carnoy’s solution 3:1 (absolute ethanol, glacial acetic acid) and stored in ethanol 70%. Different anthers and stigma positions were observed and illustrated. Photos have been taken under the binocular magnifier (x2, x4) which allows individual identification of flowers.

### Pollen viability test

The mature flower buds were bagged for the pollen viability analysis. On the day of anthesis, the flowers were collected and dissected to recover the anthers in the laboratory. The viable pollens were estimated by placing a single anther on a glass slide and crushed followed by staining with hematoxylin and eosin solution for 15 min. Finally, the pollen grain was collected and counted under a microscope. Pollen grains with bright red stains were categorized as a viable, pink stain as semi-sterile and unstained as sterile (Prasad et al., 2006). Pollen germination *in vitro*

During the bloom March-April 2018, branches with flowers in the mature stage were collected for each tree. Pollens have been released from the anthers and sown in Petri dishes containing a solidified germination medium containing 1% agar, 20% sucrose, 200 ppm of Calcium Chloride (CaCl_2_) and 75 ppm of Boric acid (H_3_BO_3_), and then incubated in the dark at 30° C for 20 h (Benlahbil and Bani-Aameur, 2002). A pollen grain was considered germinated when the pollen tube was longer than the length of the pollen grain. The observations were carried out under a microscope coupled to a digital camera for illustration.

### Fruit set

The number of fruit set was taken at the beginning of the fruit maturation stage. So, the number of fertilized flowers was calculated one month after pollination. The initial and final fruit setting was followed directly after the end of the crosses. Therefore, the set for all types of pollination was recorded 30 days after the end of the flowering and expressed as a percentage to deduce the most compatible genotypes. Intra and intercompatibility were determined as a percentage of fruit set and the number of fruits developed after successful fertilization in response to self and cross-pollination.

### Data analysis

All statistical analyses were carried out using Statistica V.12 software package. The following parameters were evaluated for variables: mean, minimum value, maximum value, standard deviation (SD) and coefficient of variation (CV %).

## Results

### Agro-morphological characterization

The descriptive analysis of the morphological data was presented in Table 1. The overall average values for the height of the trees vary from 2.67 to 5.20 m, with a minimum value of 2.60 observed for INRA-142 (pollinizer) and INRA-107 (receiver tree). All trees monitored in this study have an upright shape except for INRA-75 which has a semi-weeping shape with a very broad leaf, typical spindle fruit and very easy to break. The number of trunks per tree is relatively similar, except for INRA-132 with three trunks.

The collar diameter of 13 studied trees varied according to the number of ramifications, which ranged from 1 to 3 per tree, and the circumference presented a range from 9,33 to 16.66 m.

### Flowering periods

This study showed that there are several types of argane trees according to their flowering periods (Figure 3, Figure 4). Argane trees that bloom once a year (INRA-139, INRA-109, INRA-98, INRA-79, INRA-65 INRA- 49), argane trees which bloom twice a year over a more or less extensive period overlapping between March-May and June-July (INRA-135, INRA-132, INRA-107 and INRA-54) and argane trees that bloom throughout the year according to the three periods mentioned in Figure 4 (INRA-75).

The floral polymorphisms and the different morphs in the argane tree were studied. So, different positions of the anthers and stigma have been observed. Three morphs existing in the argane tree: Mesostylous flower has a style of intermediate length and long and short stamina, longistylous flower has long style and intermediate and short stamina and their brevistylous has a short style and stamens of intermediate and long length (Figure 5).

**Fig. 5.**
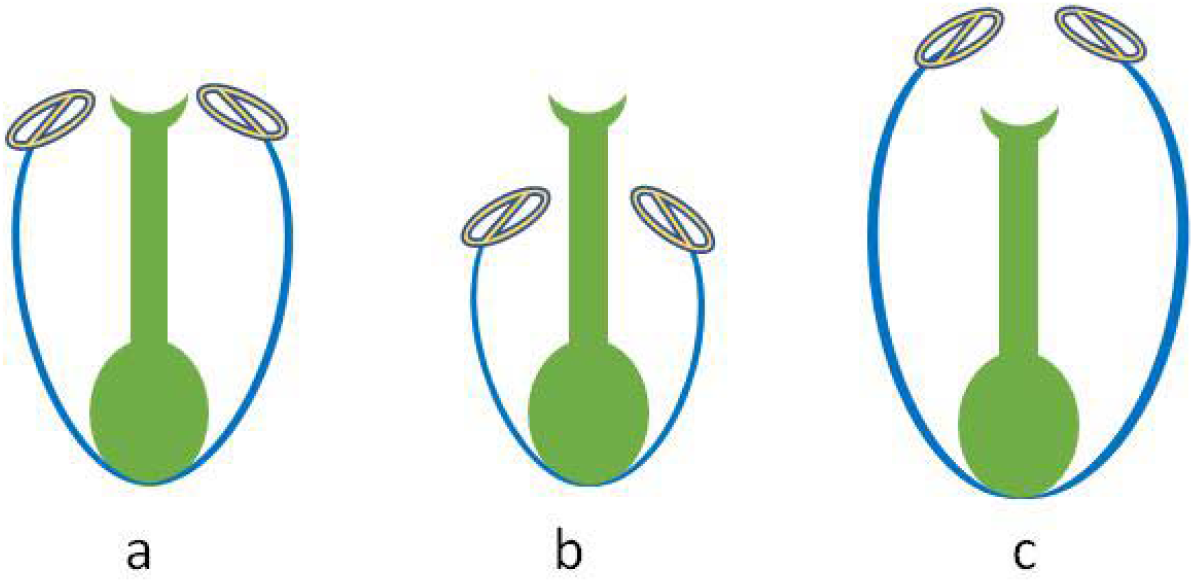
The heterostyly observed in the argane tree. (**a**): Mesostylous (**b**): Longistylous and (**c**): Brevistylous flower.

The argane female reproductive system is made up of a super gynaecium with a conical style ending in a stigma. The pubescent and superior ovary is surmounted by a short and conical style, also protruding from the stamina or vice versa (Figure 5). The ovary is made of 2 to 5 welded and uniovulate carpels. The 5 carpel gynaecium was rare. The ova are tiny and located on the central column of the gynaeceum. The placentation is of the axile type. The floral formula is: (5S) + [(5P) +5St+5E à (10E)]+ (1C) à (5C) (**Fig. 6**).

**Fig. 6.**
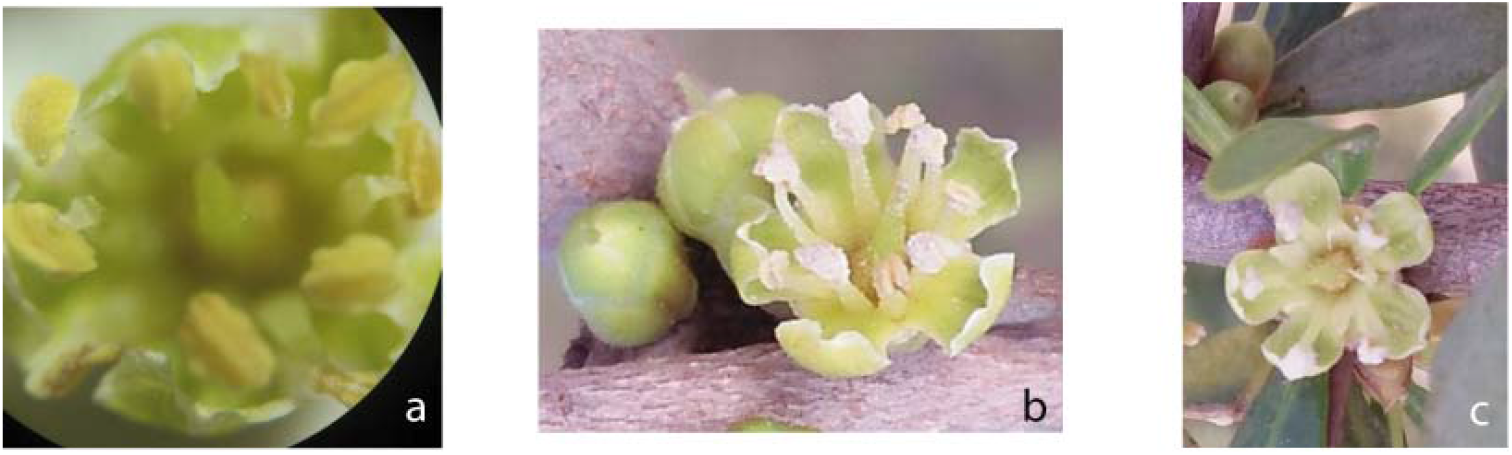
Floral morphology of argane with a flower of 10 (a), 8 (b) and 5 (c) anthers.

### Viability and *in vitro* pollen germination

The viability of the pollen assessed by staining technique showed that the pollens of trees crossed were viable and able to germinate. The pollen germination rate and the development of the pollen tube are important for fertilization. The germination of pollens on an agar medium is illustrated in Figure 7.

**Fig. 7.**
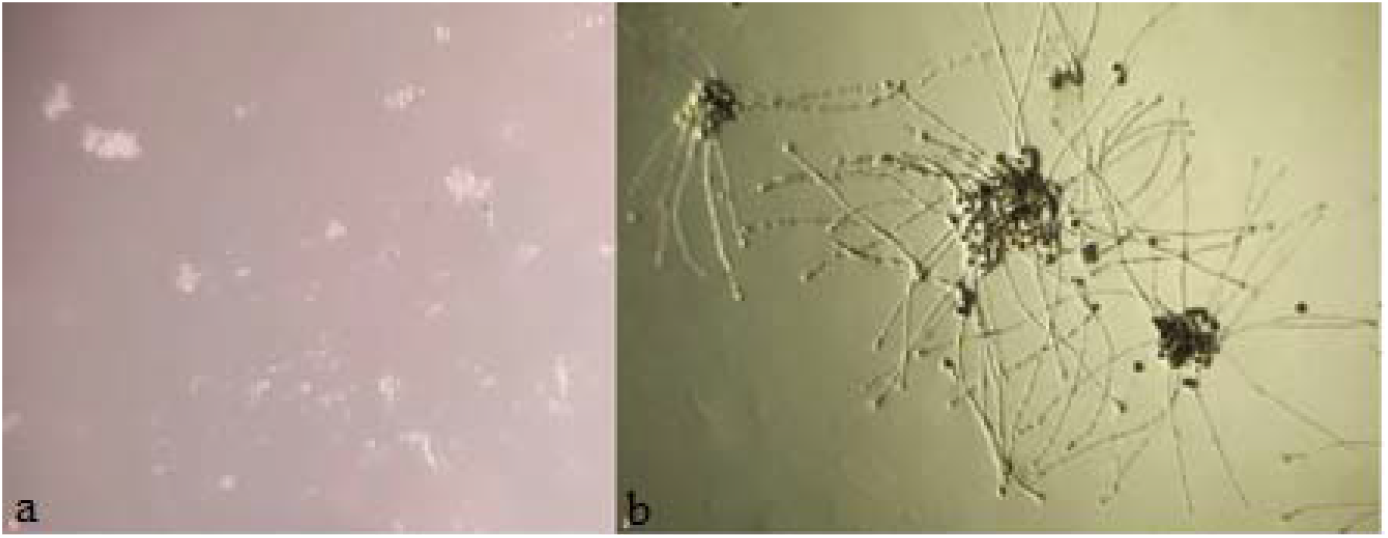
In vitro pollen germination test, (a) pollen germination with tube growth on a germination medium (1% agar and 20% sucrose), (b) absence of development of the pollen tube germinated.

### Cross combinations study

To study the effect of pollinizer on ‘female parents’ production and the crossbreeding diallel program were used on two groups during the flowering season March-April of 2018 and 2019. The 1st group is composed of 7 genotypes crossed with INRA-62 pollinizer and the second group is composed of 6 genotypes crossed with INRA-142 pollinizer (Table 2). In total, 13 cross combinations were focused in this study. In Table 2 a more detailed overview of the cross combinations along pollinizers and receiver of relative fruit set following controlled self-and cross-pollination. So, to determine the nature of the breeding system, the index of self-incompatibility (ISI) was calculated and the results show that the ISI of 11 receiver parents was less than 0 and all trees were self-incompatible (Table 2). Also, cross combinations between pollinizers and these 11 receiver parents were very efficient. The cross-compatibility rate often depends on cross associations it varies from 38.71 % to 84.38 %. Five combinations have an accounting of less than 50 % for pollinizer INRA-62 and all combinations or crossing with pollinizer INRA-142 showed that pollinizer compatibility varied from 50 to 84 %.

**Table 2.** Fruit set of cross-pollination with 11 argane genotypes.

| Pollinizers | Receiver genotypes | floral morphs | Flowers number |  | Fruits number |  | Rate of fruit set |  | Nature of breeding system |
| --- | --- | --- | --- | --- | --- | --- | --- | --- | --- |
|  |  |  | Cross/p. | Self*/p. | Cross/p. | Self/p. | Cross/p. | Self/p. |  |
| INRA-62 | INRA-139 | Mst | 66 | 115 | 30 | 0 | 45.45 | 0 | SI |
|  | INRA-135 | Bst | 62 | 9 | 24 | 0 | 38.71 | 0 | SI |
|  | INRA-109 | Bst | 14 | 168 | 8 | 0 | 57.14 | 0 | SI |
|  | INRA-79 | Mst | 36 | 15 | 24 | 0 | 66.67 | 0 | SI |
|  | INRA-65 | Mst | 79 | 64 | 39 | 1 | 49.37 | 1.56 | SI |
|  | INRA-49 | Mst | 47 | 107 | 21 | 0 | 44.68 | 0 | SI |
|  | INRA-98 | Bst | 101 | 109 | 50 | 0 | 49.50 | 0 | SI |
| INRA-142 | INRA-132 | Bst | 49 | 107 | 34 | 0 | 69.39 | 0 | SI |
|  | INRA-107 | Mst | 19 | 139 | 10 | 0 | 52.63 | 0 | SI |
|  | INRA-98 | Bst | 31 | 109 | 25 | 0 | 80.65 | 0 | SI |
|  | INRA-75 | Mst | 35 | 37 | 25 | 0 | 71.43 | 0 | SI |
|  | INRA-65 | Mst | 64 | 24 | 54 | 0 | 84.38 | 0 | SI |
|  | INRA-54 | Bst | 73 | 127 | 38 | 2 | 52.05 | 1.57 | SI |

(\*): Number of flower self-pollinated (female parent), p: pollination, SI: self-incompatible, Mst: mesostylous, Bst: brevistylous.

### Fruit set and metaxenic pollen effect on the fruit size

Although, for most cross combinations fruits are obtained at the first, but low fruit number has arrived and given the seeds mature fully. The number of fruit sets differs from one genotype to another and the response of trees via the same pollen differs from tree to tree, sometimes there is a high expression of the pollen flow, sometimes depression via other mechanisms. The presence of trees with high bloom potential and compatibility in the orchard is very effective for argane tree’s productivity. An investigation to study the effects of pollinizers and to select the best pollinizers (pollen source) were examined. Whereas, the results in most of the pollination combinations established (56 combinations from complete diallel cross programs and 6 combinations from incomplete diallel cross programs), the differences between fruit size were observed from hand-controlled pollination combinations with pollen of different sources (Figure 8). Hence, in 13 combinations that interested this study (Table 2), the difference was highly significant in fruit size between pollinizers and cross-pollinated female parents and open pollination. When we compared the fruit from cross-pollination in each combination (♂INRA-142*INRA-98♀), (♂INRA-142*INRA-65♀), (♂INRA-62*INRA-98♀), (♂INRA-62*INRA-65♀) (**Fig. 8**).

**Fig. 8.**
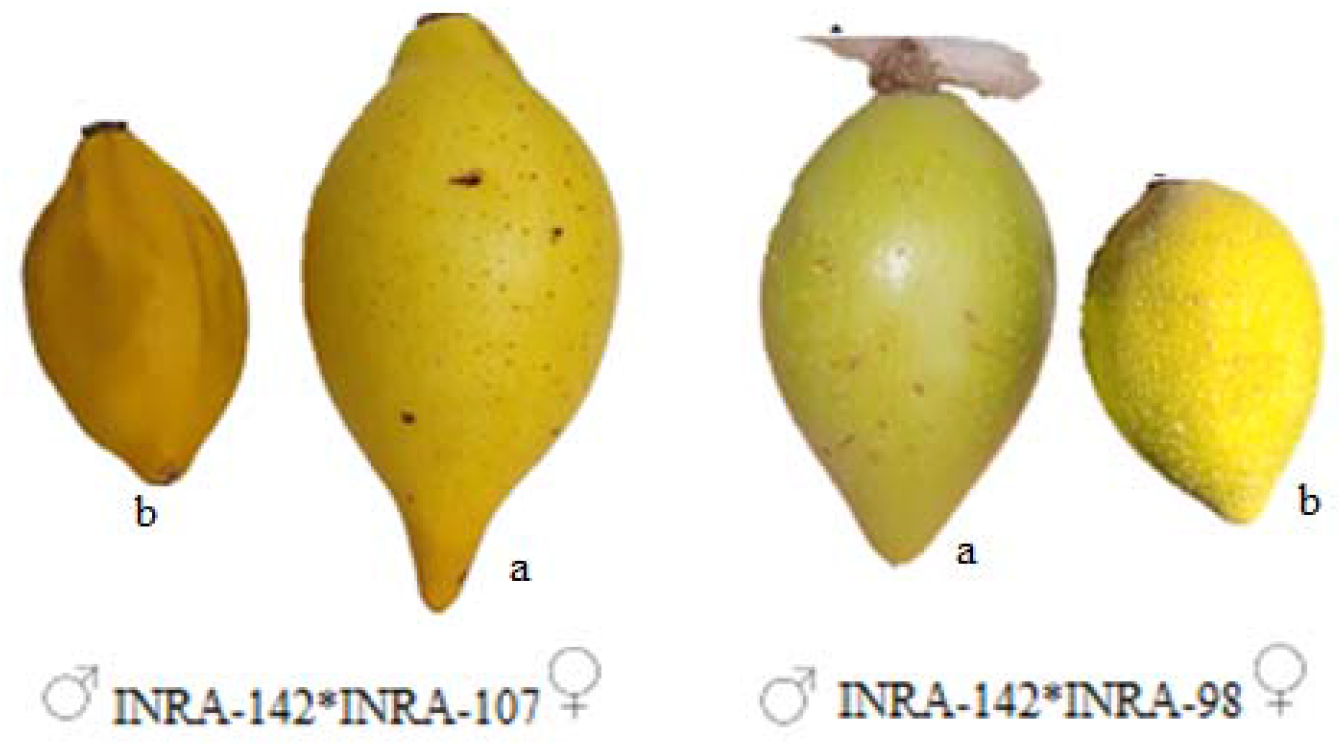
A careful choice of compatible pollinizers significantly improves the size of fruits in argane tree, (a): fruit from cross-pollination (♂INRA-142*♀INRA-98), (b): fruit from open pollination.

After, to study the floral polymorphisms and to detect the different morphs existing in the argane tree, we conducted hand pollination within and between floral morphs and examined the fruit production. A summary of the observations of fruit-set rate data calculated inter- and intra-morph hand-pollination treatments of two pollinizers (mesostylous floral morph, Mst) with 11 female parents (mesostylous and brevistylous floral morph Bst). Therefore, crossing mesostylous floral morph (floral morph style of pollinizer) with the different floral morphs (7 mesostylous and 6 brevistylous morphs floral) was investigated in the present study (Table 2). Although, the hand-pollination between (Mst)-pollinizers × (Mst)-female parents and (Mst)-pollinizers × (Bst)- female parents might be mainly caused by high compatibility between parents and high rate of fruit-set has been found for (MstxMst) (44.68–84.38) and (MstxBst) (38.71–80.65) (Figure 9, Table 2). This study shows the variations in heights of stigmata and anthers and the unequal heights of the sexual organs between floral types do not influence the fruit-set (Table 3).

**Fig. 9.**
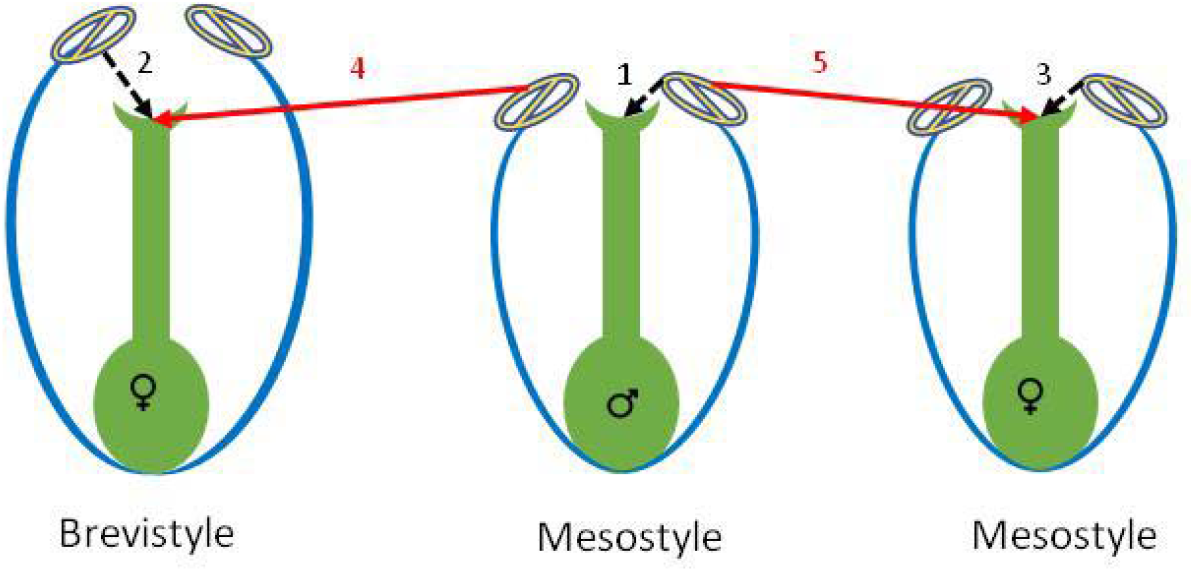
Intra and Inter-morph cross of argane tree (*A. spinosa*), Self-intra-morph crossings (dotted lines, 1-3), cross-intrer-morph (Mesostyle * Mesostyle) and (Mesostyle * Brevityle) (solid lines, 4 and 5), pollination treatments among the eleven argane trees (pollen receiver with two morphs dominated mesostyle and brevistyle) with two pollinizers of *A. spinosa* (pollen donor with mesostylemorph; pollen transfer within the same morphs but different individuals)

**Table 3.** General linear model (GLM) of the effect of pollinizers and floral morphs on fruit weight after crossings.

| variance source | Degrees of freedom | Mean square | p-value |
| --- | --- | --- | --- |
| Effect of pollinizer | 1 | 2.661 | <0.004** |
| Effect of floral morphs | 1 | 0.001 | 0.910<sup>ns</sup> |
| Pollinizer x floral morphs | 1 | 0.002 | 0,732<sup>ns</sup> |
| Error | 7 |  |  |

\* Significant; \*\* highly significant; <sup>ns</sup> not significant

## Discussion

Currently, there are scarce data regarding the pollen quality and the pollen tube growth rate during pollination (self-pollination and cross-pollination). Effective pollination is a prerequisite for fruit-set then, having information on pollen biology (viability and tube growth), is necessary for a rational approach to improve productivity and breeding programs (Bolat and Pirlak,1999; Cruzan, 1990; Shivanna, 2003). Pollen viability can be assessed by staining techniques and by pollination germination *in-vitro* and *in-vivo* tests. The choice of method depends on the species (Dafni and Firmage, 2000; Dafni et al., 2005). Several staining methods have been evaluated to distinguish between fresh and dead pollen. In the current study, after the dissection of the anthers, the pollens released were stained with hematoxylin and eosin. As a result, fresh pollens are colored red and dead pollens are not colored. Observations on the viability of pollens from crossbreed trees show that their pollens are viable and able to germinate and fertilize.

The pollen germination rate and the development of the pollen tube are important for fertilization. Thus, a low pollen germination rate can cause fruit set failure due to degradation of the ovum before the pollen tube reaches the ovary (Mellenthin et al., 1972; Therios et al., 1985). Pollen germination and growth of the pollen tube through the pistil to the ovule are regulated by pollen-pistil interactions (Linskens, 1986). The growth of the pollen tube through the pistil style has been used to study the compatibility relationships in many fruit crops, including apricots (Burgos et al., 1993; Egea and Burgos, 1996), plums (Nikolić and Milatović, 2010), almonds (Ben-Njima and Sociasi, 1995; Ortega et al., 2002; Dicenta et al., 2008), sour cherries (Cerović and Ružić, 1992) and olive tree (Selak et al., 2011).

The argane tree is a monoecious species, with hermaphrodite flowers (Perrot, 1907), alone or grouped in glomeruli appear at the base of the leaves or on the nodes of mature branches (Perrot, 1907; Bani-Aameur, 2002). Each glomerulus can contain up to 15 flowers. The evolution of these flowers takes place in six phenological phases: flower bud (FB) from 1 to 2 mm, style flower bud (SFB) from 1 to 2 mm, bloom flower (BF) from 2 to 3.2 mm, a dried flower with persistent corolla (FSCP) whose size is similar to that of bloom flower and finally the flower without corolla (FSC) from 1 to 2 mm. The full development of BF takes place in four phenological phases: BF1, BF 2, BF 3 and BF 4 (Benlahbil, 2003). The approximate lifetimes were, 45 min, 20 min, 1 hour and more than 24 hours for BF1, BF 2, BF 3 and BF 4, respectively. The lifespan of BF is at least two days. The calyx is made up of 5 free sepals (dialysepal) and a corolla with 5 free petals (dialypetal) (Bani-Aameur, 2000).

The majority of flowering plants show polymorphisms in floral morphology. Thus, heterostyly is a unique form of polymorphism. They have two (distyly) or three (tristyly) different morphological types of flowers, named “morphs” with differences in the arrangement of stigma and anther heights in the floral morphs (Barrett, 1992; Richards and Barrett, 1992). For *A. spinosa*, tristylous polymorphism with three flower forms (Mesostylous, longistylous and brevistylous) was observed as for other wild species (Nettancourt, 2001). However, for *A. spinosa* differences in style length and anther positioning influenced the interaction between the pollen and the pistil then the pollination compatibility. The morph phenotype is genetically linked to genes responsible for a unique system of self-incompatibility. Indeed, the pollen from a flower on one morph cannot fertilize another flower of the same morph. These findings of polymorphisms in floral morphology factors play an important role in the compatibility mechanism of argane tree pollination.

To choose the suitable pollinizers, in the foreground, hand pollination was conducted within and between different floral morphs and examined the fruit production. Therefore, inter-morph, cross in intra-morph (a cross between two flowers of the same morph from different individuals) and self-intra-morph (pollinated with pollen of the same individuals) pollination treatments were performed. In contrast, these observations suggest that heteromorphic compatibility shouldn’t be contributed to the failure of fruit production in inter-morph crossings and fruit-set shouldn’t be influenced by pollinizer’s flower morphs. Consequently, differences in morphological structures at stigma and style based on the relative positions of stamina and pistils do not affect fruit production for hand pollination treatments. The efficiency of insect-mediated pollen transfer in natural pollination among the different argane floral morphs is not clear and needs further studies. Whereas, less or no fruit-set of the intramorph crossing the pollinizers and female parents Bst-♂ (pollen receiver) and Mst-♀ (pollen donor) were produced. Besides, results of self-pollination treatments and the observations of pollen-tube growth^14^ suggest that pollen germinated at the surface of stigma where the pollen tubes push into style and were not blocked at the base of the stigma or further down in the style. During crossing experiments, we have observed that there is no failure of adhesion or desiccation of pollen grains on stigma (corolla still attached 48 hours after pollination) but the failure of fruit-set occurred one month after pollination. The part of postzygotic differentiation results in the inability to produce zygotes after self-pollination and the pollen tube might be inhibited before reaching the ovary, hence did not enter micropyles of any ovules. The causes of reduced pollen tubes inside *A. spinosa* style are not clear and need further studies. Also, the result of hand self-pollination confirmed that *A. spinosa* is selfincompatibles and the self-incompatibility is liable to the failure of the fruit-set. In other words, when self-pollination is first induced, the cross pollens were inhibited, but no fruit set occurred.

The semi-compatible and completely compatible argane trees need the compatible pollinizer tree for the appropriate fruit-set. Therefore, pollinizer should be compatible with female parents and have synchronization in flowering periods. Based on the results of controlled pollination, it can be concluded that argane trees are considered self-incompatible and cross-pollination is very important. Vulnerable pollination is a limiting factor in plant production in various regions (Alizadeh et al., 2009). Thus, choosing a particular pollinizer for particular female parents is currently a new topic of research for argane fruit production under arganiculture programs. Currently, there is a scarce of data regarding the effect of genetically different pollen sources on argane fruit (size or weight, color, biochemical composition…). This study aimed to evaluate the effect of two pollinizers (pollen source) INRA-142 and INRA-62 on fruit size and weight and their ability to affect the phenotype of directly developed fruits from the pollinated flowers, more precisely, these characters can present metaxenia effect in the case of fruits. This phenomenon was the first time to be evaluated for argane tree. Moreover, a comparison of the fruit from open pollination and cross-pollination showed a decrease or increase of the same characters for example the fruit size, fruit shape, color and the number of almonds per seed. Although, this observation depends on the source of pollen for the INRA-142 and INRA-62 pollinizers. While, the shape of the fruit and the maturation cycle are controlled by female parents, knowing that the argane trees differ from each other for this cycle to collect ripe fruits (from 9 to 15 months).

Based on the findings of the current study, it can be concluded that the genetically different pollen sources affected the weight and size of the argane fruit, shape, color, developmental timing and depend on the genetic background of the mother or female parents. The differential effects of fertilization with various pollen sources on the size, shape, color, or chemical composition of seeds and fruits is defined as xenia effect (Denney, 1992). The term “xenia” (first description by (Focke, 1881) is applied to direct pollen effects and covers all direct pollen effects, whether discerned in embryo, endosperm, or maternal plant tissues. However, we find another term expressed in the maternal tissues that have also been termed “metaxenia” (Swingle, 1982). Indeed, the confusion between “xenia” and “metaxenia” persists, as evidenced by the kinds of definitions available for the two words. Both xenia and metaxenia have been mainly suggested for improving plant yield. For scientists who study “xenia”, this term is used to describe direct pollen effects on (embryo and) endosperm and “metaxenia” for the effects on maternal-plant tissue. Although, “metaxenia” is applied to fruit in which fleshy carpel and accessory tissues are more economically important. Xenia is expressed in endosperm (Denney, 1992). The influence of pollen on the maternal tissue of fruits has been shown in many plants including fruit trees and several horticultural crops (Riazi et al., 1995; Sanchez et al., 2012; Rezazadeh et al., 2013; Steven et al., 2019).

This recent study, with four different pollination treatments applied to 11 argane genotypes revealed the differences between open-pollination and cross-pollination for most fruit traits (size, color, shape, time of fruit maturity), suggest the influence of the pollinizer (effect of source pollen) on fruit size and female parent effects in shape, time of fruit ripe. More researches are needed to identify suitable maternal/pollen-parent combinations that will realize some of the potentials of argane pollen parents to increase productivity. Consequently, there will further investigation in argane tree after the growth of the descendants from different crossings. Therefore, we conclude that there is a maternal control effect on the shape and time of fruit maturity and there is also an influence of the pollinizers on the size or weight of fruit and xenia reported in the argane tree for the first time. The selection of the pollen source has the potential to modify and improve argane production. This study showed that the effect of pollen-parent (xenia) occurs in all fruit components of argane tree. It is detected for the first time the occurrence of both compatible pollinizers and xenic effects of pollen on argane fruit.

## Conclusion

Synchronization of flowering is a major factor in the hybridization potential in the argane tree. The availability of genotypes with high flowering potential (pollinizer) and compatibility is very sought after, taking into account also good climatic conditions and the presence of vectors for good pollination. This study has shown that at an agronomic level, artificial pollination is feasible for successful crosses directed at the argane tree and the plantation of pollinizer in the orchard could be valuable to improve yield and quality. Identification of compatible genotypes provides an important baseline for successfully transferring traits of interest from selected elite genotypes to Argane cropping system. The controlled pollination technique showed during this study will serve as the basis for the future breeding programs of the argane tree and fruit productivity. These results from diallel crosses could be also useful in the screening of the vigorous parent with desired traits to be improved for orchard planning argane farming.

## Acknowledgements

This research was funded by the National Institute of Agricultural Research (INRA-Morocco) within the framework of the argane scientific research program. The authors would like to thank Bouzoubaa Z. for collecting and installation of plant materials and Boujghagh M. for providing expert technical support.

## Author Contributions

A.N; Investigation, Methodology, Software, Writing-article Preparation, Writing - Review & Editing; T.A; B. R; M.A Review & Editing. All authors reviewed and approved the final manuscript.

## Conflict of interest

Authors declare that they have no conflict of interest.

This article does not contain any studies with human participants or animals performed by any of the authors.

